# High impact AMPAkines induce a Gq-protein coupled endoplasmic calcium release in cortical neurons: a possible mechanism for explaining the toxicity of high impact AMPAkines

**DOI:** 10.1101/2024.01.28.577446

**Authors:** DP Radin, S Zhong, Rok Cerne, Jeffrey Witkin, A Lippa

## Abstract

Positive allosteric modulators of the AMPA receptor (AMPAkines) have a multitude of promising therapeutic properties. The pharmaceutical development of high impact AMPAkines has however been limited by the appearance of calcium-dependent neuronal toxicity and convulsions in vivo. Such toxicity is not observed at exceptionally high concentrations of low impact ampakines. Since most AMPAR are somewhat impermeable to calcium, the current study sought to examine the extent to which different mechanisms contribute to the rise in intracellular calcium in the presence of high impact ampakines. In the presence of AMPA alone, cytosolic calcium elevation is shown to be sodium-dependent. In the presence of high impact AMPAkines such as cyclothiazide (CTZ) or CX614, however, AMPAR potentiation also activates an additional mechanism that induces calcium release from endoplasmic reticular (ER) stores. The pathway that connects AMPAR to the ER system involves a Gq-protein, phospholipase C_β_–mediated inositol triphosphate (InsP3) formation and ultimately stimulation of InsP3-receptors located on the ER. The same linkage was not observed using high concentrations of the low impact AMPAkines, CX516 (Ampalex) and CX717. We also demonstrate that CX614 produces neuronal hyper-excitability at therapeutic doses while the newer generation low impact AMPAkine CX1739 is safe at exceedingly high doses. While earlier studies have demonstrated a functional linkage between AMPAR and G-proteins, this report demonstrates that in the presence of high impact AMPAkines, AMPAR also couple to a Gq-protein, which triggers a secondary calcium release from the ER and provides insight into the disparate actions of high and low impact AMPAkines.

## Introduction

Glutamate activates several neuronal ionotropic and metabotropic receptors (for review see [1]). Whereas ionotropic receptors act by fluxing ions through a channel in the receptor complex, metabotropic receptors function by stimulating intracellular second messenger systems, often via a G-protein coupling. Three types of ionotropic glutamate receptors have been characterized: α-amino-3-hydroxy-5-methyl-4-isoxazole propionic acid receptors (AMPAR), kainate receptors and N-methyl-D-aspartate receptors (NMDAR). These receptors differ in their affinity for synthetic agonists and in their gating kinetics. At resting potential, NMDAR channels are blocked by a magnesium ion, which prevents ion flux even in the presence of glutamate. This magnesium blockade is only relieved during depolarization and so while NMDAR influence the duration of fast EPSCs, they cannot induce such events. Conversely, agonist occupation of AMPAR initiates action potentials, but they deactivate rapidly due to the low affinity for glutamate and its fast release (off rate) [2].

Long-term potentiation (LTP), a phenomenon thought to be a component of memory encoding [3], has been reported to require both AMPAR activation and an elevation of post-synaptic cytosolic calcium levels (Ca^2+^_cyt_) [4]. However, most AMPAR tetramers contain Glur2 subunits that have undergone glutamine to arginine RNA codon editing rendering these receptors calcium impermeant [5]. The predominant model suggests that AMPAR facilitate LTP by inducing sodium influx and during the resulting depolarization, NMDAR and other calcium channels induce calcium entry [6]. Such events may not represent the only mechanism by which AMPAR activation elevates Ca^2+^_cyt_, since AMPAR activation even stimulates Ca^2+^_cyt_ elevation in non-excitable neural cells. In astrocytes for example, AMPAR activity elevates Ca^2+^_cyt_ by promoting the reversal of sodium / calcium exchangers (NCX) [7]. In forward mode, plasma membrane NCX expel intracellular calcium by electrogenic exchange for extracellular sodium, but during AMPAR activity they reverse direction and elevate Ca^2+^_cyt_ by excluding sodium ions [8].

Previously, it has been shown that in addition to acting as ionotropes, AMPARs also possess a G-protein linkage [9-11], indicating that they also might play a metabotropic role. The effect of positive allosteric modulators, termed AMPAkines [3], on the AMPAR metabotropic linkage is unknown, although Smith *et al*, (2000) reported that in the presence of the AMPAkine cyclothiazide (CTZ), AMPAR activation of astrocytes depletes endoplasmic reticular (ER) calcium stores. This CTZ-dependent connection between AMPAR and ER stores was not further elucidated but might be evidence for an AMPAkine-dependent metabotropic AMPAR pathway.

A wide variety of AMPAkines have been reported and, despite being derived from different chemical structures, all share the common property of enhancing the actions of glutamate at these receptors (for review see Alt et al [12]). However, using electrophysiological and receptor binding approaches, Arai and colleagues [3, 13, 14]. have described two different categories of AMPAkines based upon differences in the way these classes modulate AMPAR. Type I or high impact AMPAkines, such as CTZ, CX546 and CX614, enhance agonist binding affinity and interfere with agonist-induced AMPAR desensitization. In contrast, Type II or low impact AMPAkines such as aniracetam, CX516 and CX717 do not modulate agonist binding and have little effect on agonist-induced AMPAR desensitization. Rather, these low impact AMPAkines accelerate ion channel opening and slow deactivation. While high impact AMPAkines have shown disease modifying activities in a number of CNS pathologies, they seem to produce neuronal toxicity [15], an effect not seen with low impact ampakines. This delineation warrants a close examination as to why high but not low impact ampakines elicit neuronal toxicity.

The present study will examine the mechanism(s) by which AMPAR activation elevates Ca^2+^_cyt_ in the presence and absence of representative high and low impact AMPAkines. This study will examine the role of sodium in agonist-dependent responses and the AMPAkine-dependent effect of AMPAR stimulation on intracellular calcium release. A pathway connecting AMPAR stimulation to ER calcium release will be elucidated and an AMPAkine-dependent G-protein linkage will be described that is only active in the presence of high impact AMPAkines. As a corollary, the seizurogenic activity of high impact AMPAkines, but not low impact AMPAkines, will be expounded upon which may be explained by the selective linkage to ER calcium stores by high impact AMPAkines.

## Materials and Methods

### Solution preparation

Solutions were prepared by addition of drug in dimethyl sulfoxide (less than 0.1 % of final volume) to a siliconized microcentrifuge tube. Controlled salt solution (CSS: 125 mM NaCl, 25 mM glucose, 25 mM HEPES, 5.0 mM KCl, 1.4 mM CaCl_2_, 0.8 mM Mg_2_SO_4_) was added and the drug was dispersed by vortex mixing. For sodium-free experiments, NaCl was replaced with 110 mM N-methyl-D-glucamine (NMDG, Sigma)and 15 mM Tris-base. The CSS used in manganese studies contained only 100 μM CaCl_2_instead of 1.4 mM. When solutions of lipophilic drugs such as U73122 (Tocris) were prepared, a high-power sonic probe was used to ensure dispersal.

### Neuronal cultures

Primary cortical neuron cultures were prepared according to standard methods with slight modifications [16]. Briefly, the cortices and hippocampi were dissected from E15-17 Wistar rats. The tissue was cut into small pieces and incubated for 9 min with 1 mg/ml papain in Ca^2+^ and Mg^2+^ free Hank’s Balanced Salt Solution (HBSS) at 37 °C. After this time, the solution was replaced with high glucose Dulbeco’s modified Eagles medium containing 10% fetal bovine serum (FBS), 2 mM glutamine, 100 IU penicillin and 100 μg/ml streptomycin (DMEM). The tissue was dissociated by repeated passage through a pipette after addition of 50 ug/ml DNAse. The resultant cell suspension was centrifuged through a 5% bovine serum albumin solution and the pellet was re-suspended in DMEM. The cells were counted and seeded at a density of 3800 per mm^2^ onto the inner 60 wells of a poly-D/L-lysine coated black-sided 96 well plate (Corning #3603) in a final volume of 200 μl DMEM per well. The outer wells of the plate were filled with water to reduce evaporation and the cultures were maintained at 36 °C and 7% CO_2_ for 7-10 days before use.

### Cytosolic calcium concentration (Ca^2+^_cyt_) measurements

After 7 to 10 DIV, the culture media was replaced with 100 μl of 2 μM Fluo-4-AM in CSS. The cells were incubated for 15 min at room temperature, after which time the dye was replaced with 100 μl fresh CSS. The cells were then allowed to sit undisturbed for 5-10 min before an additional 100 μl of CSS containing AMPAkines plus or minus investigational drugs was carefully added. Cultures were then allowed to stand for 45 min prior to initiation of the study. Changes in Ca^2+^_cyt_ were quantified by recording the fluctuation in light emission at 520 nm caused by excitation of the Fluo4-dye by a 485 nm light beam (Fluostar Galaxy fluorimeter, BMG Labs, USA). After taking six baseline fluorescence values at 0.1 s intervals, studies were initiated by adding 10 μl of 100 μM (s)-AMPA (final concentration = 5μM). Changes in fluorescence were then recorded for 6 s in 0.1s intervals.

### Ca^2+^_cyt_ measurements in the presence of Manganese

The protocol used for the examination of intracellular calcium release was a modification of the procedure published by Merritt et al [17]. The procedure was similar to that described above except the CaCl_2_ concentration was reduced to 100μM and 10μl of 3mM MnCl_2_ (final concentration= 150 μM) was injected at t=-10s. AMPA was injected at t=0 and changes in *Ca*^*2+*^_*cyt*_ fluorescence were recorded at 0.2s intervals for 20s following AMPA addition.

### Inositol phosphate (InsP) Assay

Intracellular InsP levels following AMPAR stimulation were assessed using cortical neurons plated in the inner wells of 24 well plates (Biotrak [3H]IP3). After 10 DIV the medium in the dishes was replaced with 0.2ml of CSS containing 100 μM Ca^2+^. 10 min later 0.1ml of CSS or CSS containing 1.2mM CX614 or 120 μM CTZ was added. 0.1 ml of CSS containing 40 mM lithium and Group I mGluR antagonists; 2-Methyl-6-(phenylethynyl)pyridine hydrochloride (400 nM MPEP) and 7 (Hydroxyimino)-cyclopropachromen-1a-carboxylate ethyl ester (160 μM CCPCoEt) was also added. Study was initiated by addition of 25μl of a CSS containing 1.7mM EGTA and either 85 μM AMPA or 170 μM NBQX. After 30 min of stimulation the CSS was removed, and cells lysed by addition of 0.5ml ice-cold 5% perchloric acid. Twenty minutes later, the acid was neutralized with 1.5M KOH, universal indicator and 10mM Tris. The supernatant was then transferred to microfuge tubes, the pH adjusted by color to approximately pH 8.0 and then the samples were taken to dryness under vacuum. To analyze samples were resuspended in 500 μl of distilled water, 750 ul of anionic exchange resin (Dowex AG 1X-8; formate) was added. After centrifugation, the resin was washed with water and inositol phosphates were eluted in 600μl of 1.5mM NH_4_COOH : 0.1mM HCOOH and 300 μl of each sample was subjected to scintillation counting.

### Extracellular recording in hippocampal slices

Standard techniques were used to prepare transverse hippocampal slices (400 μm) from 130 – 200 g male Sprague-Dawley rats. After dissection, slices were placed in an interface chamber and perfused at 0.75 ml/min with ACSF (124 mM NaCl, 26 mM NaHCO3, 10 mM glucose, 3.4 mM CaCl_2_, 3 mM KCl, 2.5 mM MgSO_4_, 1.25 mM KH_2_PO_4_) saturated with 95% O_2_ / 5% CO_2_. Field EPSPs were recorded in the *stratum radiatum* of region CA1 in response to activation of the Schaffer-commissural fibers by a stimulating electrode located between the recording sites. Stimulating pulses of 0.05-0.1 ms were delivered every 20 s using a paired pulse protocol with an interpulse interval of 200 ms. Evoked potentials were band-pass filtered using frequency limits of 0.1 Hz and 10 kHz and recorded using Neurodata Acquisition Software (Theta Burst Corp, CA).

To determine the maximum EPSP amplitude achievable without spiking, the input –output relation of the synaptic response was assessed, and the stimulation intensity adjusted to produce response amplitudes that were about 50% of the maximum. After calibration, the slices were allowed to stabilize, and the experiment only commenced after 20-30 min of stable baseline recordings.

### Fluorescence data analysis

Data were analyzed by two different techniques depending on the model used. When examining extracellular sources of calcium influx, data was analyzed by examining the area under the curve (AUC_t=0.5-5s_) resulting from plotting the change in fluorescence of each well over time. Fluorescence level in each well was assessed for several seconds prior to AMPA addition in order to calculate mean baseline values. When examining intracellular calcium release (the Mn^2+^ assay) however, an apparent “ceiling” in Ca^2+^_cyt_levels limited the usefulness of AUC analyses. Under such conditions linear regression of the slope was found to yield more representative data (as indicated by lower inter-replicate variability and improved “fit” of values to sigmoidal concentration response curves). Linear regression was performed by examining the slope of the linear part of the fluorescence curve that occurred between 2 and 18 s (usually 3-12 s) after AMPA injection. These times were chosen empirically because the first 2-3 s after AMPA addition acted as a “lag phase” and responses tended to lose linearity over time as the “ceiling” was approached.

### Excised membrane patch-clamp recording

Outside-out membrane patches were obtained from CA1 pyramidal neurons present in 400 μm cultured hippocampal slices taken from 13-day-old Sprague-Dawley rats. The patch pipettes were filled with 65 mM CsF, 65 mM CsCl, 10 mM EGTA, 2.0 mM MgCl_2_, 2.0 mM ATP disodium salt and 10 mM HEPES (pH 7.3). The pipette tip was immersed in a solution of 125 mM NaCl, 2.5 mM KCl, 1.25 mM KH_2_PO_4_, 2.0 mM CaCl_2_ 1.0 mM MgCl_2_, 5.0 mM NaHCO_3_, 25 mM glucose and 20 mM HEPES (pH 7.3). The patch was exposed to 1mM glutamate alone or with selected concentrations of CX717 or CX614 using a switch-flow technique described previously [18].

### Slice Seizure Assay

Extracellular field potentials recording (Population Spikes) in hippocampal slice CA1 was used to determine potential seizure liability. Transverse 300-400 μm hippocampal slices were taken from adult male Sprague-Dawley rats (4–6 week-old) and incubated in a submerged slice recording chamber system for 30 to 60 min. The chamber was perfused with oxygenated aCSF as described above at room temperature. Population Spike (PS) recordings were made using glass electrodes filled with 2M NaCl (1-5megaohm) placed in the region of pyramidal cell body of CA1. Electrical stimuli were delivered with twisted bipolar nichrome wire electrodes in the Schaffer-commissural fiber afferents. Population spike was induced by activating Schaffer-commissural fiber using single pulse stimulation delivered at 0.05 Hz. The maximal amplitude of PS was determined by increase stimulating intensity until a second spike appeared. Then the intensity was decreased to induce a 50% to 60% of maximal response. The amplitude of PS was measured for each response and plotted against time. At the same time, the amplitude of second PS and/or number of PS was measured and plotted against time. The size of the second PS and numbers of PS were used to estimate the seizure potential or seizure state. The perfusion then was switched to control ACSF and continued to record until drug effect was washed back to the baseline. Compounds were prepared by dissolving in DMSO and then added to ACSF.

### Statistical Analysis

In fluorescence studies, experiments consisted of 5 replicate wells; the responses from which were averaged to generate a mean value. All experiments were performed on 3 to 5 different occasions and at least two different neuron preparations. Figures show the mean values of a representative study plotted as means ± SEM. Tabular values and histograms represent the mean values of several studies (n=15-40), wherein each replicate is expressed as a percentage of the mean positive control for that experiment and then data are presented as a mean value ± SEM. Statistical significance was generally examined using ANOVA together with a post-hoc Dunnett multiple comparison test, although in the hippocampal slice studies or IP3 formation assays, a matched pair, one-tail t-test was used. In all cases, significance was attributed when pΣ0.05 (*) or pΣ 0.01 (**). The data were analyzed and plotted using the software packages “Prism” (GraphPad, CA) and “Microsoft Office” (Microsoft, WA).

## Results

Addition of AMPA to cortical neuron cultures stained with Fluo-4 (a calcium sensitive fluorescent dye) resulted in a rapid (<0.5s) increase in cytosolic calcium (Ca^2+^_cyt_) that reached a horizontal asymptote within 1-2s of AMPA addition and was antagonized by the addition of the AMPA antagonist 10 μM NBQX (Fig 1a). The effect was maximal at 5 μM AMPA with an EC_50_ of 1.5 μM (Fig 1b). The maximal Ca^2+^_cyt_ level achievable with AMPA alone was increased further in wells that contained a high impact AMPAkine. Figures 1c and d show a secondary, slower onset time-response curves from cortical cultures stimulated with 5 μM AMPA in the presence of CX614 or cyclothiazide (CTZ), two structurally distinct high impact AMPAkines. This dose-dependent response was observed over several seconds, in contrast to the rapid response elicited by AMPA treatment alone. Both of the two low impact AMPAkines, CX717 or CX516 only minimally affected the rapid, initial phase of AMPA-induced Ca^2+^_cyt_ elevations and the second late phase, even at mM concentrations (Fig s1).

**Figure 1:**
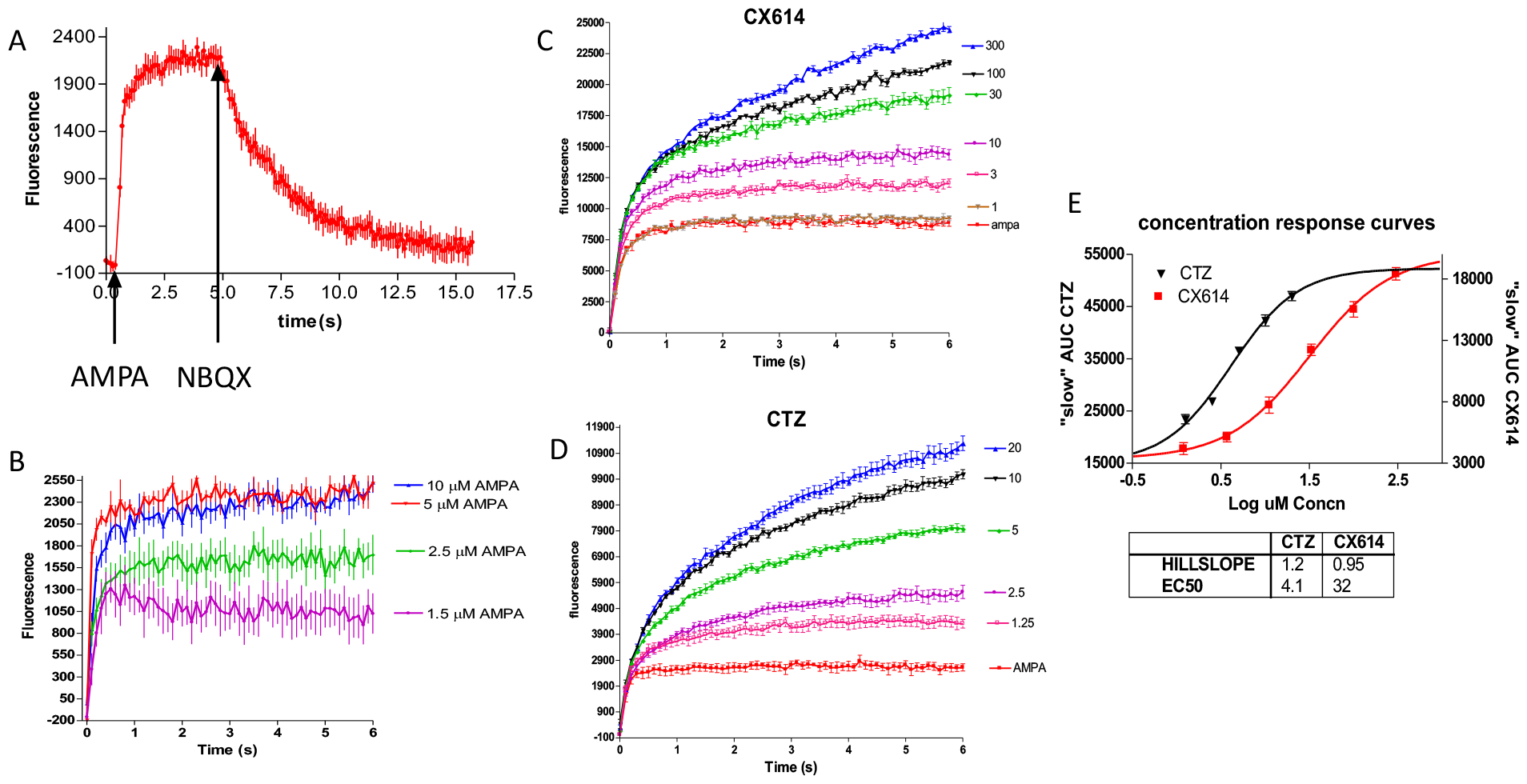
Injection of AMPA +/-AMPAkines induces varying degrees of calcium elevation in Fluo-4 stained cortical cultures, as measured by increase in emitted light at 520 nM. Baseline fluorescence was subtracted from responses. A) AMPA increases in fluorescence are amenable to AMPA antagonist NBQX (n=5 wells). B) Calcium increases with concentration of AMPA. Maximal response was observed with 5 μM AMPA (n=5 wells). C & D) The “slow” component of the calcium elevation response to AMPA addition that occurs in the presence of C) CX614 or D) CTZ is dependent on ampakine concentration. E) AUC_t=0.5-5s_ values for calcium responses following AMPA injection plotted against log ampakine concentration.

Responses induced in the presence of AMPA plus a high impact AMPAkine include a concentration-dependent rising sustained “slope”, with the E_max_ for CX614 and CTZ occurring at 300uM and 20uM, respectively. This is in contrast to the significantly lower asymptotic level achieved by any concentration of AMPA alone (Fig 1a). AMPAkine-enhanced slope responses were quantified by calculating areas under the curve between t=0.5-5s (AUC_t0.5-5_). Non-linear fit analyses of the data showed EC_50_ values for CX614 and CTZ to be 32 and 4.2 μM respectively (Fig 1e). To determine whether this effect might be due to high impact AMPAkine-induced interference with desensitization [3], responses produced by AMPA with and without a high impact AMPAkine were compared with those induced by kainate (80μM), a slowly-desensitizing AMPAR agonist. Kainate-induced responses were comparable to those seen with AMPA in the absence of a high impact AMPAkine, with no slope response observed (data not shown).

In order to determine the sodium dependence of the Ca^2+^_cyt_ rise, sodium in the culture medium was replaced with N-methyl-d-glucamine (NMDG), a large organic anion incapable of transiting through AMPAR. Figure 2a demonstrates that when sodium in the media was replaced with NMDG, AMPA-induced Ca^2+^_cyt_ elevations were abolished, but in the presence of a high impact AMPAkine, AMPA continued to induce Ca^2+^_cyt_elevation. Furthermore, concentration-dependent Ca^2+^_cyt_ responses to CX614 and CTZ observed in sodium-depleted media showed comparable EC_50_ values to those observed in sodium containing media (Fig 2b-d).

**Figure 2:**
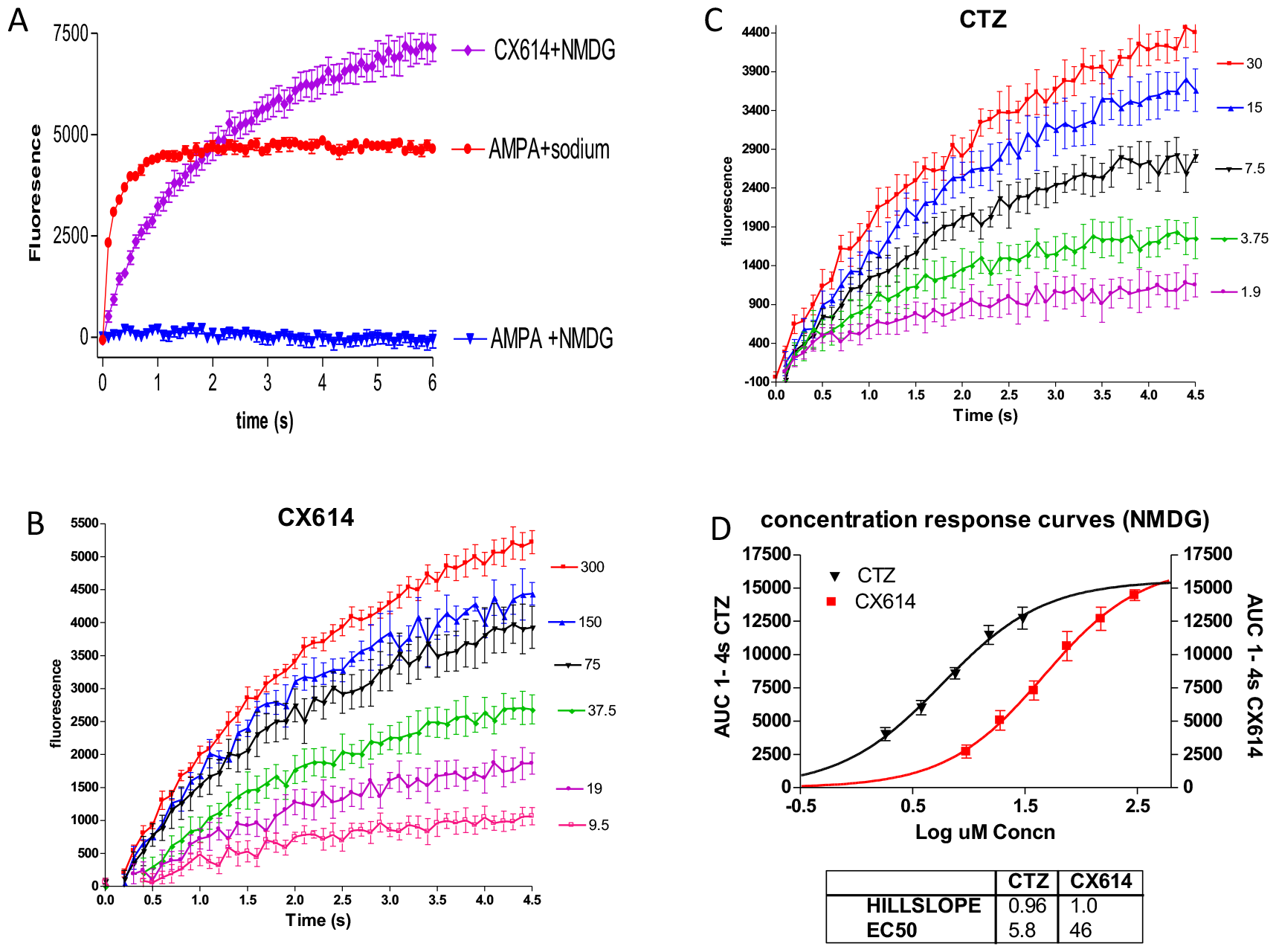
Replacement of sodium in the medium with an equimolar concentration of N-methyl-d-glucamine (NMDG) eliminated the effect of AMPA on cytosolic calcium levels, but not the ampakine-dependent “slow” component response. A) AMPA in the presence of NMDG does not induce cytosolic calcium elevations, whereas, 40uM CX614 still produces the monophasic “slow” component rise, which is quenched by NBQX addition. B) Calcium response to AMPA is dependent on CX614 concentration in the well (n=5 wells). C)Calcium response to AMPA is dependent on CTZ concentration in the well (n=5 wells). D)AUC_t=0.5-5s_ values for calcium responses following AMPA injection plotted against log ampakine concentration (n=5 wells).

High impact, but not low impact, AMPAkine-facilitated AMPA stimulation of Ca^2+^_cyt_responses was also maintained when extracellular Ca^2+^ was reduced to 100 μM Ca^2+^ (Fig S2), suggesting the possible involvement of intracellular calcium stores. However, the plate reader used in this study was not capable of resolving subcellular systems. In order to gain sufficient signal to noise to investigate intracellular calcium stores, studies were conducted in media containing reduced (100 μM) extracellular Ca^2+^ with 150 μM Mn^2+^ added 10s prior to AMPA addition. Merritt *et al* [17] reported that calcium dyes do not fluoresce when bound to Mn^2+^, an observation that was confirmed with the current dye (Fluo4). Figure 3a presents data demonstrating that Fluo-4 fluorescence is quenched by Mn^2+^ with an EC_50_ that is 7-fold lower than that of Ca^2+^. Addition of Mn^2+^ improved the signal to noise by inhibiting fluorescence caused by extracellular dye, (i.e., excreted from cells into the media) and by blocking voltage-sensitive calcium channels [19]. Figures 3 b and c show AMPA plus high impact AMPAkine responses for CX614 and CTZ conducted in the presence of 100μM Ca^2+^ and 150 μM Mn^2+^. The EC_50_values observed under these conditions (Fig 3d) were comparable to those observed in aCSF and in sodium depleted media (Figs 1d, 2d). This system together with pharmacological tools, was further used to examine the effects of AMPAR activation in the presence of high impact AMPAkines on intracellular calcium signaling.

**Figure 3:**
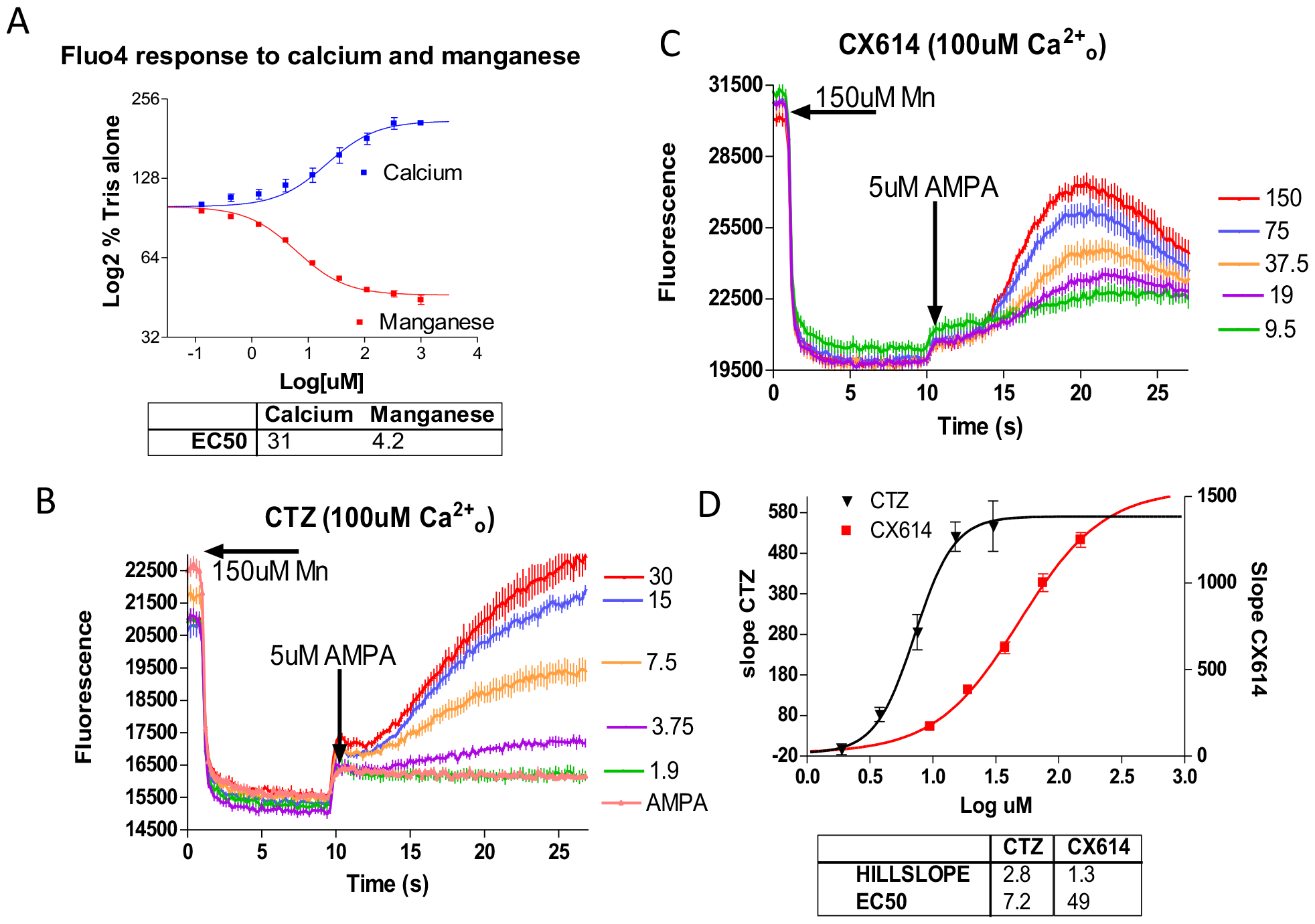
Fluorescence studies in a Mn^2+^ containing solution. A) Fluorescence of a 20 nM solution of Fluo4 (25 mM Tris, pH 7.4) at 520 nM is approximately doubled by Ca^2+^ with an EC_50_ of 31μM. Manganese on the other hand, halved the fluorescence, with an EC_50_ of 4.2 μM (n=3). B & C) Injection of manganese at t=0 reduced baseline fluorescence. After a new baseline was achieved, 5 μM AMPA was injected at t=10s. In addition to the small immediate AMPA response, a delayed and slower response was observed, dependent on B) CX614 or C) CTZ concentration (n=5 wells). D) Slope values (based on linear phase of response) for calcium responses plotted against log ampakine concentration (n=5 wells).

The results summarized in Table 1 show that AMPAR-AMPAkine-dependent responses were eliminated by pretreatment with 1μM thapsigargin, which blocks smooth endoplasmic reticulum (ER) calcium ATPases [20]. Consistent with this finding, responses were also inhibited by 75 μM 2-APB, which blocks inositol triphosphate (InsP3)-mediated ER calcium store release [21]. However, 2-APB also blocks extracellular Ca^2+^ influx through store-operated channels (SOC) [21]. Therefore, control studies were conducted in the presence of 100 μM lanthanum, which also blocks SOC [22]. Unlike 2-APB, lanthanum did not significantly block AMPAkine responses (<10±10%, p=0.92), suggesting that 2-APB inhibited AMPA-AMPAkine responses by blocking InsP3-mediated release of ER calcium stores. Furthermore, 10mM lithium, an inhibitor of InsP3 phosphatase [23], strongly potentiated (>300%) the slope of AMPA plus AMPAkine Ca^2+^_cyt_ responses. InsP3-mediated ER calcium release is often a downstream result of G-protein mediated phospholipase Cβ (PLC) activity. U73122 (10 μM), a PLC inhibitor [24], inhibited AMPA plus AMPAkine responses by more than 80% (see table 1).

**Table 1:** Involvement of Endoplasmic Reticulum in ampakine-mediated cytosolic calcium increase.

| Site of Action | % untreated ampakine response |  |  |  |  |  |  |  |
| --- | --- | --- | --- | --- | --- | --- | --- | --- |
|  | Thapsigargin |  | 2-APB |  | Lithium |  | U73122 |  |
|  | CX614 | CTZ | CX614 | CTZ | CX614 | CTZ | CX614 | CTZ |
| SERCA | 07<br>±02** | 02<br>±02** |  |  |  |  |  |  |
| IP3R/SERCA<br>/SOC |  |  | 17<br>±03** | 22<br>±03** |  |  |  |  |
| Inositol<br>phosphatase |  |  |  |  | 300<br>±150** | 350<br>±80** |  |  |
| Phospholipase C |  |  |  |  |  |  | 18<br>±03** | 13<br>±03** |

Calcium responses to 5 μM AMPA and either 100 μM CX614, 15 μM CTZ in the presence of 100 μM $\text{Ca}^{2+}$ and 150 μM $\text{Mn}^{2+}$ were inhibited by 30 min pretreatment with 1 μM thapsigargin, 75 μM 2-APB, 10 μM U73122, and strongly potentiated by 15 min pretreatment with 10 mM $\text{LiCl}_2$ . Responses were unaffected by 100 μM $\text{LaCl}_3$ . \*\* $p < 0.01$ , Dunnett multiple comparison test compared to untreated ampakine normalized fluorescence value. SERCA - Sarco/endoplasmic reticulum $\text{Ca}^{2+}$ ATPase.

One mechanism by which AMPAkine-enhanced AMPAR activation could result in InsP3-mediated intracellular Ca^2+^ release would be by stimulating release of neurotransmitters such as glutamate that then act on established metabotropic systems. To examine this possibility, the formation of inositol phosphates in neuronal cultures stimulated with AMPA in the presence of AMPAkines was examined (Fig 4) in the presence of 2mM EGTA (to prevent calcium stimulated events) and the metabotropic glutamate receptor antagonists MCPG (500 uM) or in the presence of a combination of the mGLuR 1 and 5 antagonists, CPCCOEt (40 uM) and MPEP (500 nM) (Fig 4). In both sets of studies, AMPA alone did not significantly increase InsP3 levels (compared with that produced in the presence of NBQX), although the combination of AMPA and AMPAkine significantly increased InsP3 production compared with AMPAkine and NBQX.

**Figure 4:**
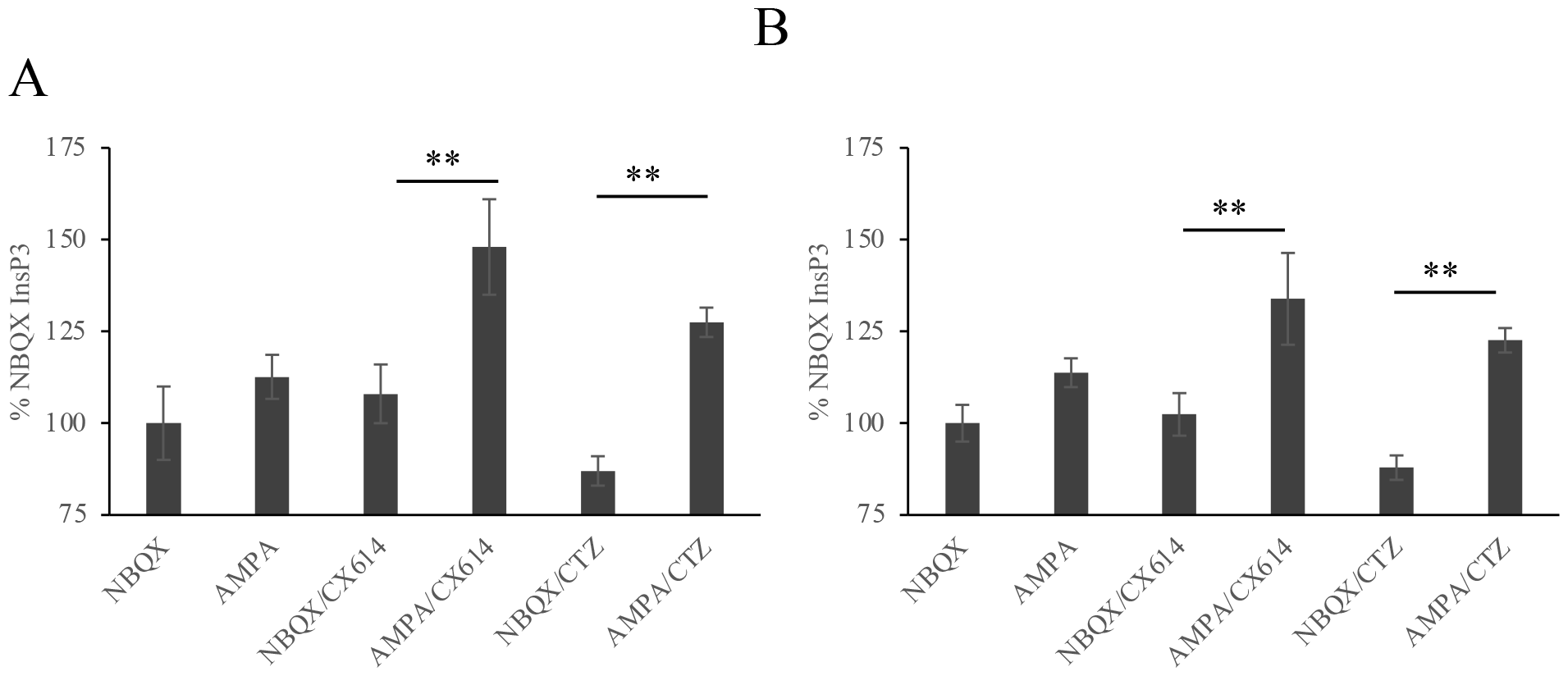
Effect of AMPA receptor positive allosteric modulation on InsP3 production. Column chromatography was used to assay for InsP3 production 30 minutes after treatment with NBQX/AMPA in the presence or absence of CX614 or CTZ. A) Cortical cultures were pre-treated with Li and EGTA to prevent InsP3 breakdown and to prevent calcium-dependent events, respectively and with MCPG, a broad metabotropic glutamate receptor antagonist. Data are normalized to cortical neurons treated only with NBQX. **p<0.01, t-test, compared to cells treated with respective ampakine with NBQX. B) Cortical cultures were pre-treated with Li, EGTA and CPCCOEt (metabotropic glutamate receptor 1 antagonist) and MPEP (metabotropic glutamate receptor 5 antagonist). Data are normalized to cortical neurons treated only with NBQX. **p<0.01, t-test, compared to cells treated with respective ampakine with NBQX. Data are displayed as average +/-SEM of 4 independent experiments.

The question of secondary neurotransmitter release was also examined in a more general fashion by examining AMPA plus AMPAkine Ca^2+^_cyt_responses in a media containing 100μM Ca^2+^ and 150 μM Mn^2+^ following 1 h pretreatment with a cocktail containing 25 nM tetanus toxin, an inhibitor of vesicular transmitter release and 200 nM tetrodotoxin, a blocker of sodium channels preventing depolarization. The cocktail did reduce responses (P<0.05) but only modestly; 18±5% in the presence of CX614, and 19±5% in the presence of CTZ (P<0.05) (Table 2). Taken together, results from these calcium and InsP3 studies support an involvement of InsP3-mediated intracellular calcium release in AMPA plus AMPAkine Ca^2+^_cyt_ responses and suggest they are most likely not a result of secondary neurotransmitter release.

**Table 2:**
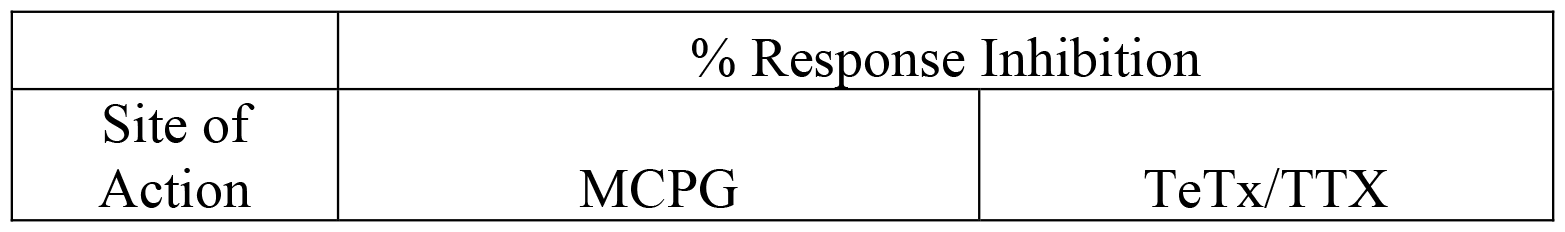
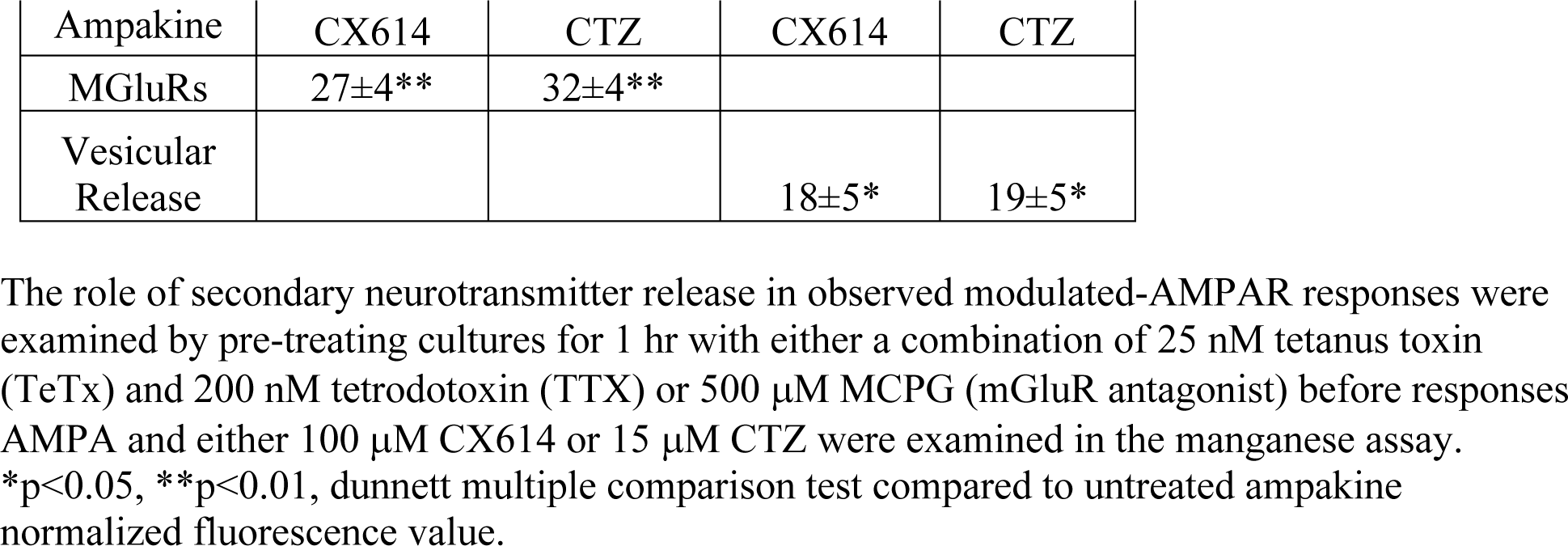
Secondary neurotransmitters exhibit minimal effects on ampakine-mediated cytosolic calcium level increases.

|  | % Response Inhibition |  |
| --- | --- | --- |
| Site of Action | MCPG | TeTx/TTX |

|  |  |  |  |  |
| --- | --- | --- | --- | --- |
| Ampakine | CX614 | CTZ | CX614 | CTZ |
| MGluRs | 27±4** | 32±4** |  |  |
| Vesicular Release |  |  | 18±5* | 19±5* |

The role of secondary neurotransmitter release in observed modulated-AMPA responses were examined by pre-treating cultures for 1 hr with either a combination of 25 nM tetanus toxin (TeTx) and 200 nM tetrodotoxin (TTX) or 500 μM MCPG (mGluR antagonist) before responses AMPA and either 100 μM CX614 or 15 μM CTZ were examined in the manganese assay.
\*p<0.05, \*\*p<0.01, dunnett multiple comparison test compared to untreated ampakine normalized fluorescence value.

As has been described before [3, 14], the effects of CX717 and CX614, on glutamate-induced currents in cultured hippocampal neurons distinguished the electrophysiological differences between these two AMPAkine classes. When administered alone, a 1-2 second application of glutamate produced a rapid depolarization followed by a rapid, prolonged desensitization with a steady state current approximately 3% of the peak level. Concurrent administration of CX614 produced a dose-dependent increase in steady-state current with an EC50 of 30uM. At maximal concentrations CX614 produced a total block of desensitization of glutamate-induced currents, with steady state current surpassing by 143 +/-18% over the peak current elicited by application with glutamate alone (Fig S3). In contrast, CX717 only increased the steady-state current to a much smaller maximum of approximately 25% of the peak response.

Further studies examined whether the AMPAkine-dependent pathway can also be observed in hippocampal slices, a model which possesses a more natural spatial relationship between neurons and supporting cells than exists in cell cultures. Acutely prepared hippocampal slices were subjected to a repetitive “paired pulse” stimulus of the Schaffer-collaterals and the resultant field excitatory post-synaptic potentials (fEPSPs) in region CA1 were recorded at 20-s intervals. In each of the three slices we observed that the amplitude and “half-width” (Fig 5; a measure of response duration) of the second fEPSP response to stimuli were both substantially enhanced by 10 μM CX614, in a manner that reversed when the drug was washed out (Fig 5a, b). Application of the InsP3 receptor antagonist, 25 μM 2-APB, at a concentration that blocked AMPA plus high impact AMPAkine responses by approximately 80% in cultures (Table 1), reduced the half-width of CX614-enhanced responses by approximately 30% (Fig 5a-c) but did not reduce CX614 enhancement of fEPSP response amplitudes. The 2-APB inhibition of CX614-enhanced fEPSP’s half-width dissipated upon 2-APB removal (Fig 5a) and 2-APB did not affect baseline fEPSP responses (data not shown).

**Figure 5:**
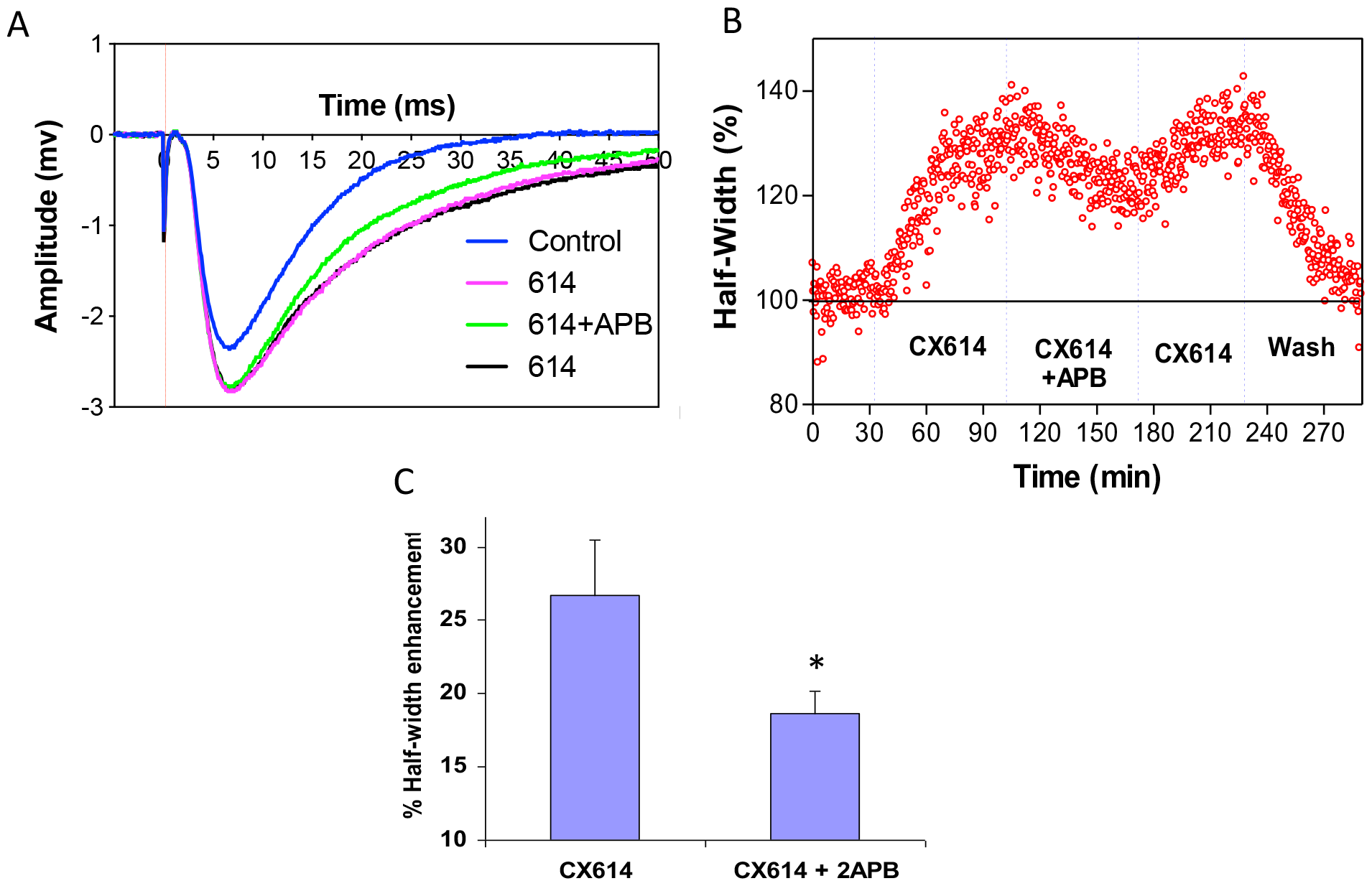
Excitatory post-synaptic potentials (EPSPs) resulting from “paired pulse” stimulation of schaffer-collaterals were recorded in CA1 at 20s intervals. CX614 increased the “half-width” (a measure of response duration) in a manner that was antagonized by 2-APB. A) Representative trace of EPSP resulting from paired pulse stimulation of Schaffer-collaterals of CA1 after addition of CX614 followed by 2-APB and subsequent application of CX614 alone. B) Typical response of EPSPs following CX614 application. 10 μM CX614 enhanced the half-width of stimulated EPSPs, while 25 μM 2-APB application during CX614 infusion antagonized CX614-enhancement. Inhibition of CX614-enhancement “washed out” upon 2-APB removal. C) Percentile inhibition of stimulated fEPSP halfwidth by 10 μM CX614 or CX614 plus 25 μM APB in CA1 hippocampal slices. (mean ± SD n=3 slices). *P<0.05, t-test, compared to half-width data in the absence of 2-APB.

Finally, we sought to determine the effects of high and low impact AMPAkines using an in vitro slice seizure assay model [25-28] (Fig. 6). Single pulsed electrical stimulation was used to evoke spikes in hippocampal slices. The spike amplitudes stabilized after 20 minutes with average spike amplitudes of 1.8 +/-0.1mV. Approximately 1 minute after 50 μM picrotoxin was perfused at the same speed as control solution, a second PS of lower amplitude appeared with the first spike remaining at the similar 1.8 mV amplitude, as in the control conditions (Fig 6b,c). Similar to picrotoxin, 16 uM CX614 drastically up modulated seizure-like electrical activity. CX614 (16 μM) produced an increase in the number of spikes per stimulation to 5-6 spikes, whereas picrotoxin only produced 1 additional population spike. In addition, CX614 increased the amplitude of the population spike. With regards to the effects of low impact AMPAkines in this assay, we utilized the newer generation low impact ampakine CX1739, which is being developed to treat opiate-induced respiratory depression and neurological sequelae secondary to spinal cord injury. CX1739 failed to modulate seizure activity in the slice preparation up to its solubility limit of 200uM (Fig s4). These data highlight an important therapeutic distinction between high- and low-impact AMPAkines, which may be related to their effects on Ca^2+^_cyt_.

**Figure 6:**
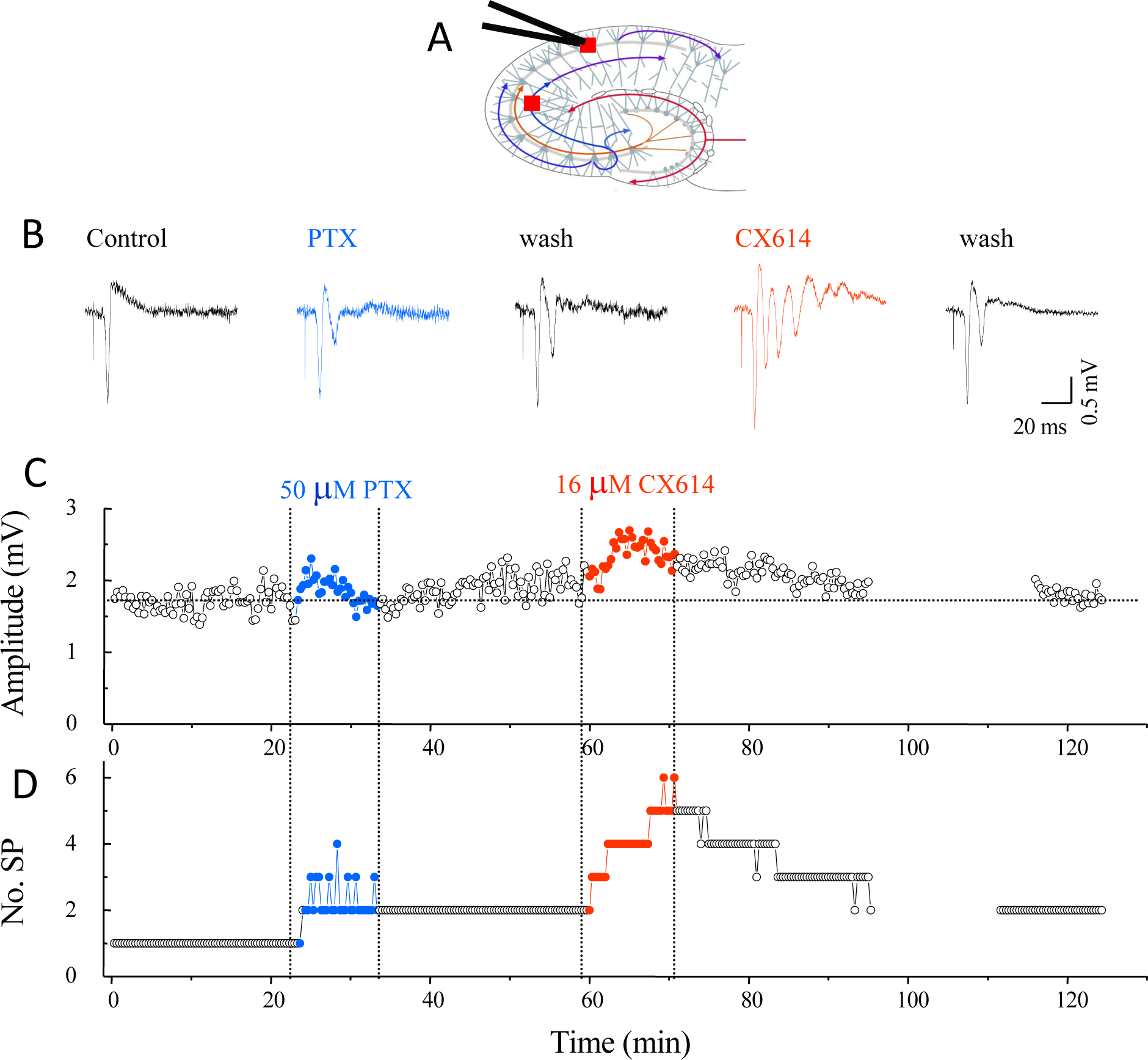
Effect of picrotoxin and CX614 on CA1 evoked population spikes in rat hippocampal slices. A) Representative schematic of recording procedure. B) Representative traces taken from periods of control and after 50 uM picrotoxin (GABA-A antagonist, positive control) or 16uM High Impact ampakine CX614 washed in CA1 pyramidal cells. C) Peak amplitude of population spikes D) Number of spikes per trace after wash in of control solution, picrotoxin, or CX614.

## Discussion

Exposure of cortical neurons to AMPA produces qualitatively and quantitatively different Ca^2+^_cyt_ elevation responses in the presence and absence of high impact AMPAkines (Fig 1). Administration of AMPA alone induced a rapid monophasic response that did not occur in sodium-depleted (hyponatric) media. In the presence of high impact AMPAkines, a second slower response was additionally observed that increased Ca^2+^_cyt_ levels above the maximal levels observed with AMPA alone. No such induction of a slow response was observed with either of the two low impact AMPAkines (CX516 and CX717). The slow response was not dependent on sodium influx (Fig 2). The magnitude of the secondary response (as determined by area under the curve) was AMPAkine concentration dependent, and two structurally distinct AMPAkines (CX614 and CTZ) exhibited EC50’s commensurate with published data of their electrophysiological effects (compare Fig 1 with [3, 29, 30]. These results indicate that the secondary response is not due to an off-target effect but is mediated by AMPARs.

The EC_50_values of slow responses induced with either of the two high impact AMPAkines were similar in normo- and hyponatric conditions. When taken together with data from cultures treated with tetrodotoxin (under hypocalcemic conditions), these results indicate that the high impact AMPAkine-dependent component is independent of membrane depolarization and may involve release of intracellular calcium stores. In order to examine this hypothesis, the fluorometric plate reader assay was modified by reducing extracellular calcium to the minimum concentration found to preserve intracellular calcium store responses (100 μM) and introducing 150 μM manganese (Mn^2+^) 10 sec prior to AMPA stimulation. This technique, a modification to that developed by Merrit et al [17], relies on calcium indicator dyes having a higher affinity for Mn^2+^ than calcium, and that the Mn^2+^complex is non-fluorescent. In the case of Fluo4, the dye used in the current study, the affinity for Mn^2+^ is seven-fold greater than for calcium (Fig 3a), so that Mn^2+^ introduction quenched baseline fluorescence. Serendipitously, Mn^2+^ also inhibits voltage sensitive calcium channels [19] and plasma membrane sodium: calcium exchangers [31], thereby also reducing extracellular calcium influx.

In the modified-Merritt assay, AMPA stimulation alone exerted only a very modest increase in Ca^2+^_cyt_, whereas responses to AMPA plus either of the high impact AMPAkines produced biphasic responses, with the slow component being proportionally much greater than in the normo-calcemic assay. The high impact AMPAkine-dependent responses continued to exhibit EC_50_values consistent with those previously observed. Studies using inhibitors aimed at the smooth endo-reticular calcium store pathway demonstrated that high impact AMPAkine responses require PLCB and smooth ER InsP3-receptors and endoreticulum ATPases. Prevention of InsP3 breakdown with lithium substantially enhanced high impact AMPAkine-dependent calcium fluorescence, but had no effect on responses induced by AMPA alone. This finding indicates that no evidence of AMPAR-InsP3-dependent responses can be observed unless a high impact AMPAkine is present, even when responses are amplified by inhibiting InsP3 breakdown.

Results similar to those observed in the current study might be expected, however, if modulated-AMPAR stimulation caused neurotransmitter release, which then activated other Gq-protein coupled receptors. This hypothesis was tested by inclusion of a broad spectrum mGluR antagonist and, in another experiment, by pretreatment with a combination of tetanus and tetrodotoxins (which prevent neurotransmitter release and neuronal depolarization, respectively). Neither treatment blocked more than a small proportion of modulated-AMPAR-responses, further strengthening the hypothesis of a direct high impact AMPAkine-dependent AMPAR-Gq-protein coupling.

To examine whether desensitization might be related to the Gq-protein coupling, an experiment was conducted using the slowly desensitizing AMPAR agonist, kainate, instead of AMPA. Another experiment was also performed to investigate whether AMPA might have an effect on InsP3 synthesis following lithium pretreatment. The data from these experiments showed that although lithium greatly enhanced AMPAkine-dependent secondary slow response, it had no discernable effect on the rapid (<0.5sec) AMPA-alone response. Similarly, stimulation with kainate did not induce a response different from that of AMPA administered alone, observations that further the conclusion that occupation of a high impact AMPAkine binding site is required for Gq-protein coupling; AMPAR-agonists alone do not induce ER calcium release. Other disparate physiological effects of kainate and AMPAkines have been shown previously, as AMPA+CTZ but not kainate induce choline release in primary neuronal cultures [32].

This study elucidates a modulated-AMPAR coupling to calcium release from the ER. This pathway’s existence also could be inferred from two previous reports. Liu et al [33] described CTZ-dependent InsP3 formation in oligodendrocytes and interpreted their data to suggest that CTZ induced a large increase in InsP3 production. However, if the data are re-examined, with AMPAR-antagonist values being used to describe baseline values (thereby eliminating effects of endogenous glutamate release), Liu’s data actually demonstrate that CTZ is *required* for InsP3 production. Similarly, in astrocytes, Smith et al [7] reported that AMPAR activation depleted thapsigargin-sensitive calcium stores, presumably from ER, in the presence of CTZ, but not with agonist alone. Together, these reports suggest that the AMPAR-ER coupling elucidated in the current study may be active in several neural cell types.

The data presented here also raise the possibility that transient response potential (TRP) channels play a role in the high impact AMPAkine-dependent slow calcium rise. The 7 subfamilies of TPV channels are very selective for their specific cations and have been reported to regulate calcium influx following Gq-protein activation. Previous reports have implicated their important role in regulating the influx of extracellular calcium into post-synaptic neurons [20]. In the modified Merritt assay, the fact that either thapsigargin or 2-APB, an IP3 receptor blocker, decreased ∼80-95% of the AMPAkine response suggests that the majority of the AMPAkine-mediated slow phase calcium rise is due to the liberation of ER calcium stores.

All of the data discussed above demonstrate the effects of modulated-AMPAR on ER calcium release in dissociated neurons, but the relevance of this pathway on the physiological actions of AMPAkines is unclear. To probe this question, an acute hippocampal slice preparation was used. In hippocampal slices, stimulation of the Schaffer collaterals evokes electrophysiological responses, the amplitude and duration (as measured by the half-width) of which are enhanced by CX614 [3]. Application of the InsP3 receptor antagonist, 2-APB, significantly inhibited CX614-induced half-width enhancement (Fig 5), without altering amplitude enhancement or affecting the amplitude or half-width of control responses. This result suggests that AMPAkine-dependent action-potential prolongation may be at least in part due to calcium release from ER stores.

In addition, the disparate seizurogenic activity between high and low impact AMPAkines (Figs. 6 and S4) and that only a modest enhancement in glutamate-induced steady state currents is required to produce a pronounced pharmacological effect, especially with regards to reversing opiate-induced respiratory depression [34-37] or alleviating respiratory depression after spinal cord injury [38-40].

In summary, stimulation of AMPAR on cortical neurons causes a sodium-dependent cytosolic calcium increase, but in the presence of high impact AMPAkines, AMPARs also couple to a Gq-protein, which results in a secondary, slower, stimulation of calcium release from the ER. This secondary effect might contribute to the prolongation of synaptic responses induced by high impact AMPAkines and might explain why high impact AMPAkines [32] but not low impact AMPAkines [41] (Table 3) have been shown to produce excitotoxic cell death in neuronal cells and convulsions in intact animals [15].

**Table 3.**
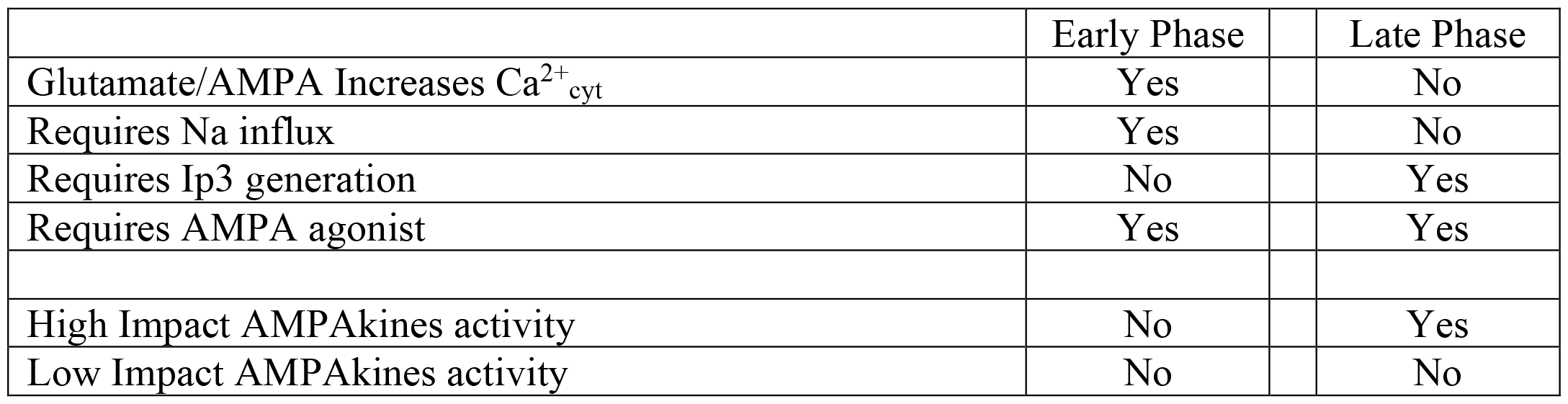
AMPAR Stimulated Increases in Ca^2+^_cyt_.

## Figure Legends

Figure s1: High doses of the low impact ampakines CX516 and CX717 do not induce a slow component of the calcium elevation response after AMPA addition (n=5 wells).

Figure s2: Extracellular calcium reduction (∼100 uM) reduces AMPA-dependent rises in cytosolic calcium levels but the slow rise remains with pre-treatment of CX614 but not CX717 (n=5 wells).

Figure s3: High and low impact AMPAkines differentially augment glutamate-induced currents in patches excised from CA1 pyramidal cells.

A) Representative traces of current induced by glutamate in the presence and absence of CX717 (n=5 wells). A dose-dependent increase in steady-state current elicited by CX717 with an EC50 of 3.4 uM.

B) Representative traces of current induced by glutamate in the presence and absence of CX614. A dose-dependent increase in steady-state current elicited by CX614 with an EC50 of 30uM.

Figure S4: Low impact AMPAkine CX1739 is not active in the hippocampal slice seizure assay.

A) Representative traces taken after 1 minute of perfusion with 0-200 uM CX1739.

B) Amplitude and number of spikes of traces elicited by perfusion of control solution or 100 to 200 uM CX1739.

