## Supplementary material for "High impact AMPAkines induce a Gq-protein coupled endoplasmic calcium release in cortical neurons: a possible mechanism for explaining the toxicity of high impact AMPAkines": 4 Supplemental figures

### Slide 1
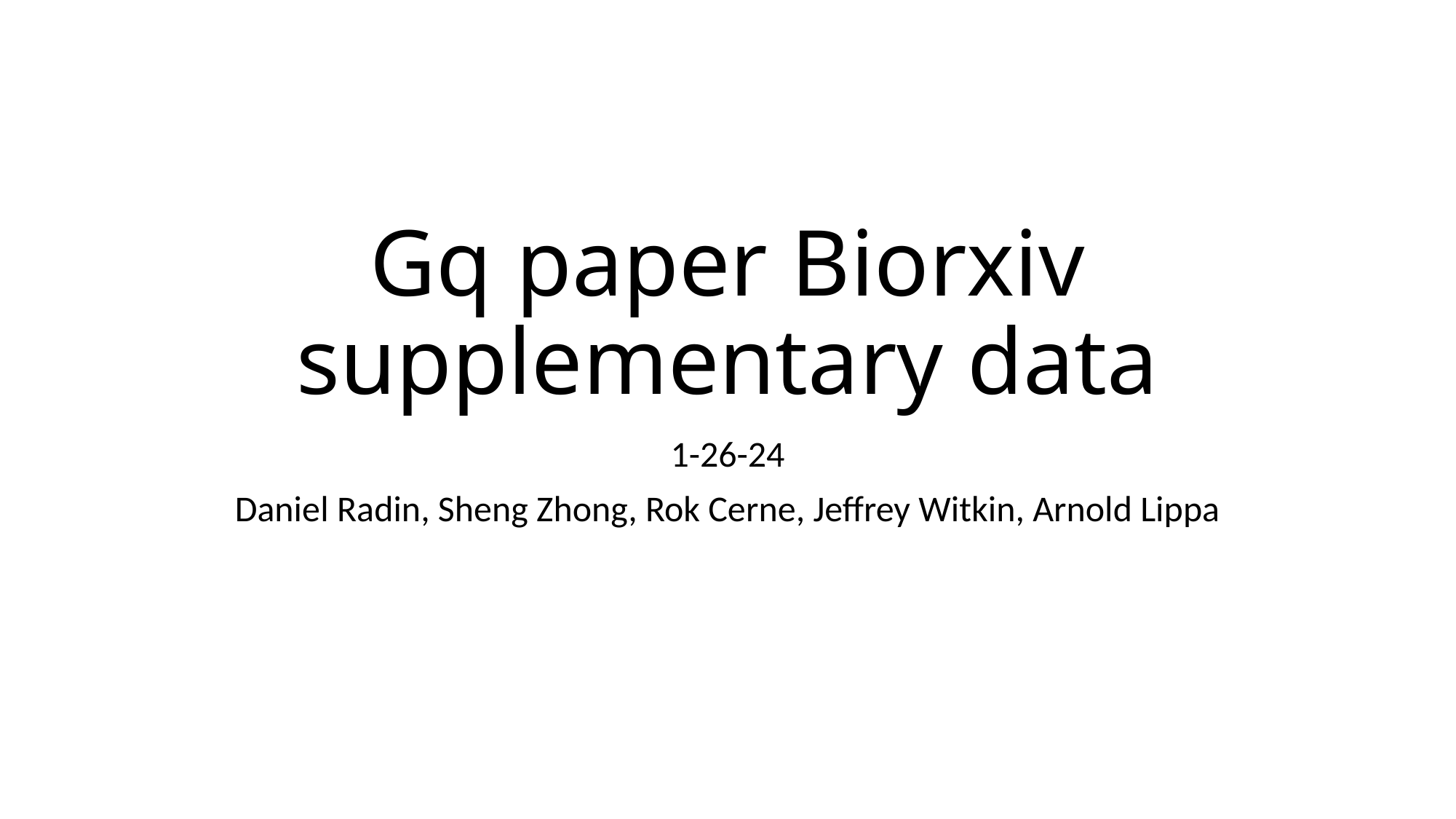

#### Gq paper Biorxiv supplementary data
1-26-24
Daniel Radin, Sheng Zhong, Rok Cerne, Jeffrey Witkin, Arnold Lippa

### Slide 2
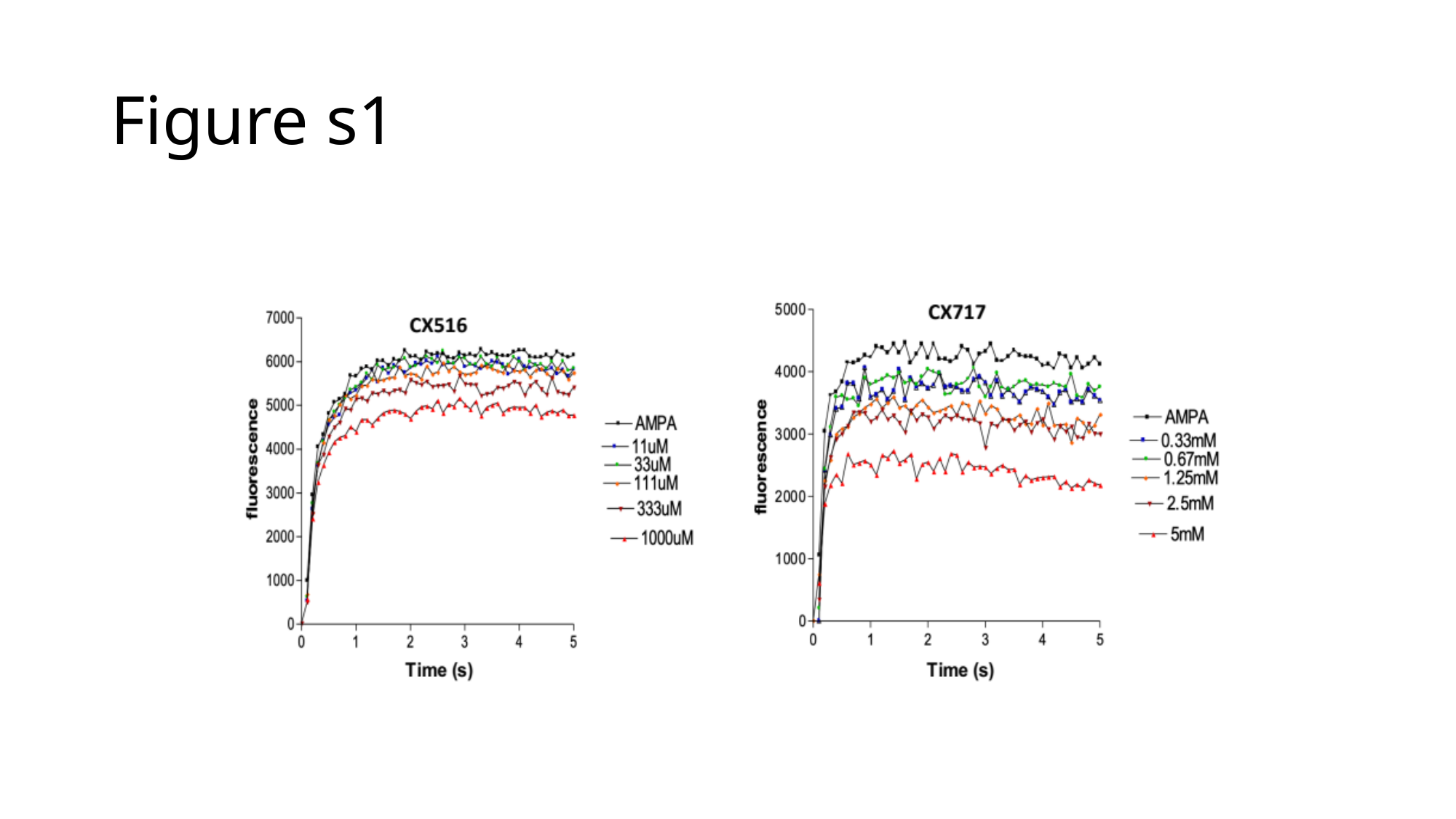

#### Figure s1

### Slide 3
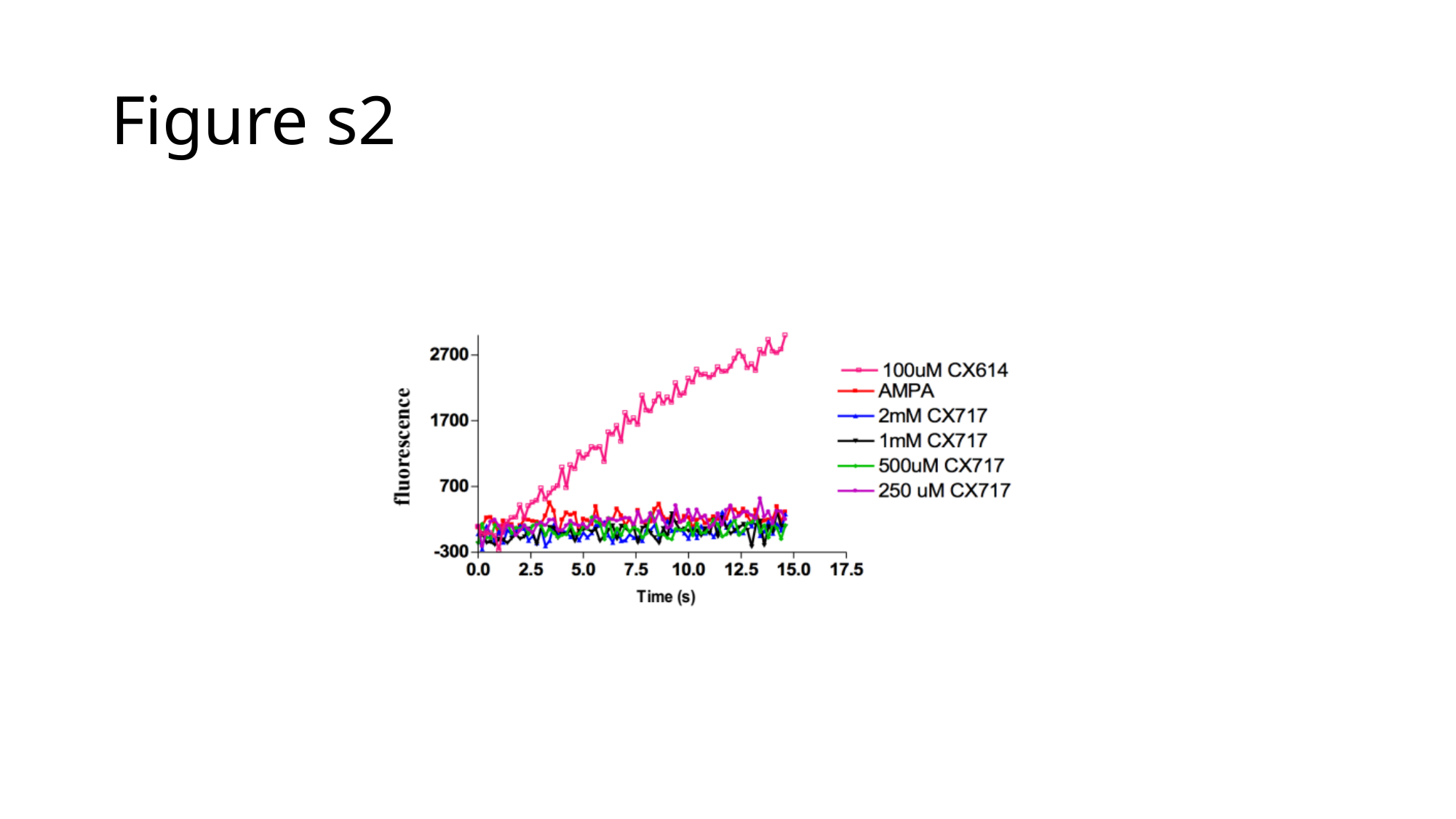

#### Figure s2

### Slide 4
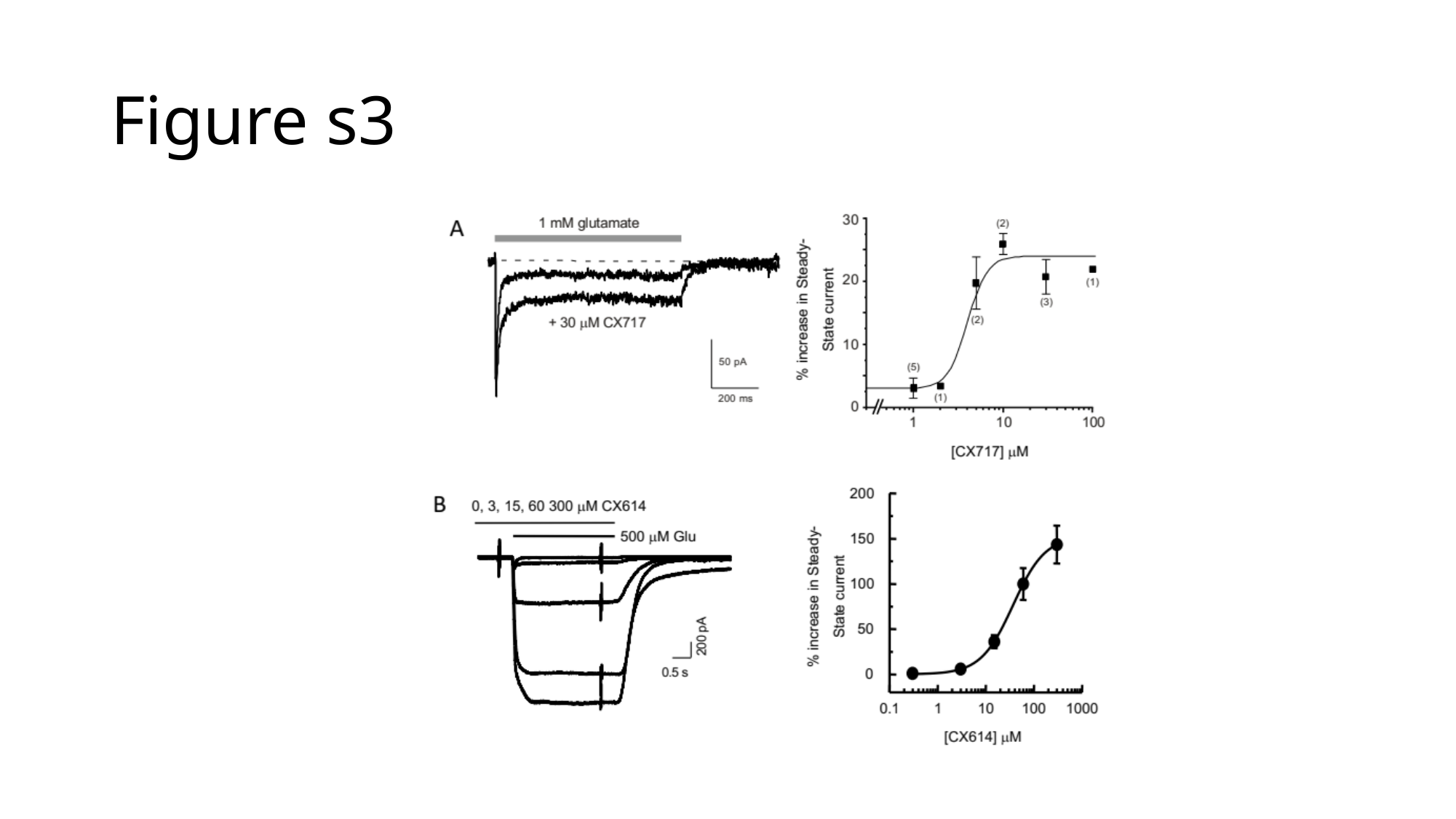

#### Figure s3

### Slide 5
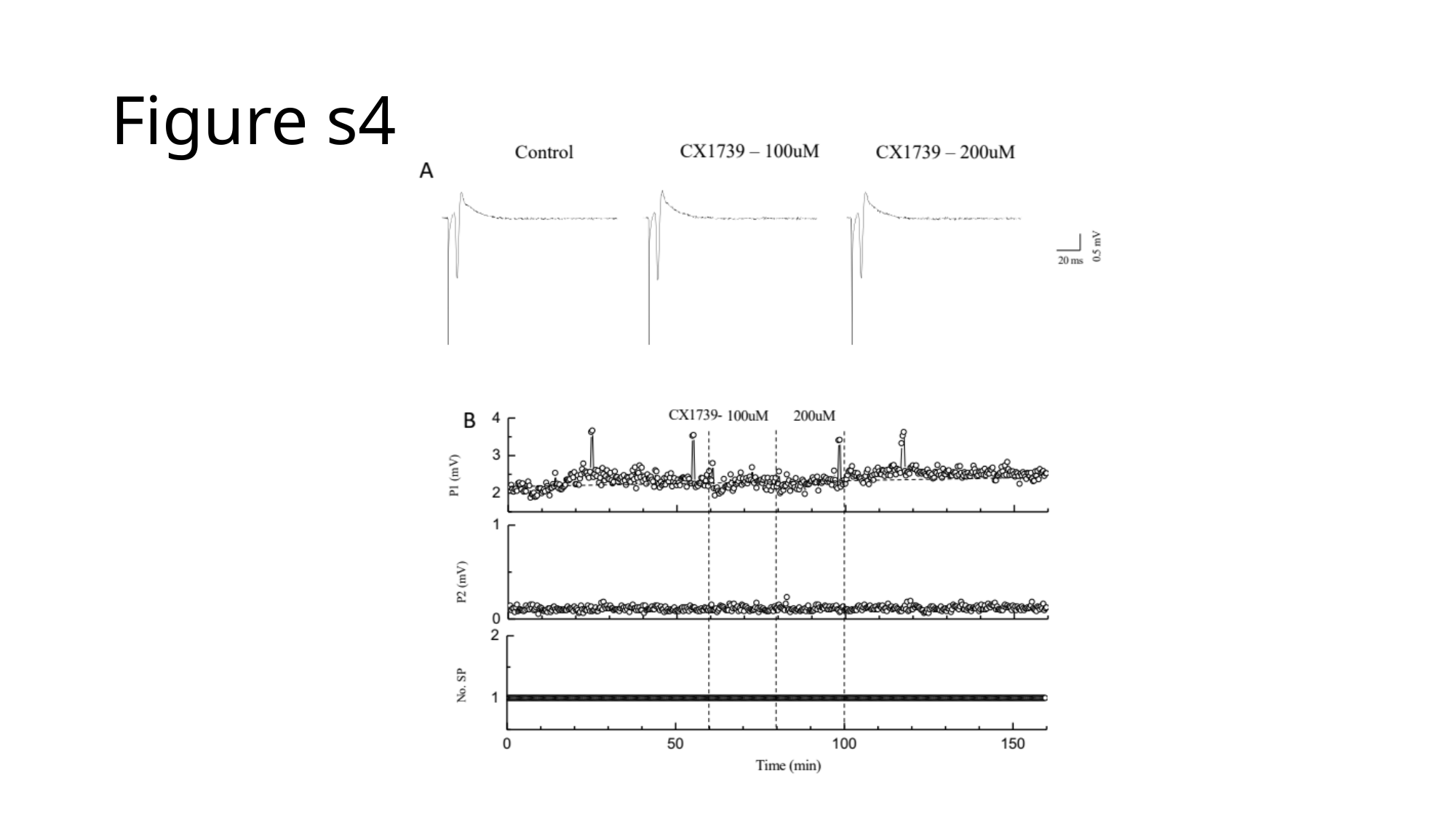

#### Figure s4
